# Elephants develop wrinkles through both form and function

**DOI:** 10.1101/2023.08.24.554618

**Authors:** Andrew K. Schulz, Noemie Reveyaz, Lena Kaufmann, Cindy Ritter, Thomas Hildebrandt, Michael Brecht

## Abstract

The trunks of elephants have prominent wrinkles from their base to the very tip. But neither the obvious differences in wrinkles between elephant species nor their development have been studied before. Asian elephants have more dorsal major, meaning deep and wide, trunk wrinkles (~126 ± 25 SD) than African elephants (~83 ± 13 SD). Both species have more dorsal than ventral major trunk wrinkles and a closer wrinkle spacing distally than proximally. In Asian elephants wrinkle density is high in the ‘trunk wrapping zone’. Wrinkle numbers on the left and right sides of the distal trunk differed as a function of trunk lateralization, with frequent bending in one direction causing wrinkle formation. MicroCT-imaging and microscopy of newborn elephants’ trunks revealed a constant thickness of the putative epidermis, whereas the putative dermis shrinks in the wrinkle troughs. During fetal development wrinkle numbers double every 20 days in an early exponential phase. Later wrinkles are added slowly, but at a faster rate in Asian than African elephants. We characterize the lifelong development of trunk wrinkles in Asian and African elephants and discuss the relation of species differences in trunk wrinkle distribution and number with behavioral, environmental, genetic, and biomechanical factors.

## Introduction

Animals have developed different mechanisms for navigating, manipulating, and interacting with their environments. These mechanisms often have clear form-function relationships where the mechanism has a form, such as a wrinkle on the elephant trunk, to satisfy a function, e.g. allowing the trunk to stretch while maintaining protection^1^. Elephant trunks are their primary tool of manipulation and interaction with their environment and are essential for sensory perception such as olfaction and somatosensation^2,3^. The elephant trunk is described as one of the three prominent examples of muscular hydrostats along with octopus arms and mammalian tongues^4^, but elephants are unique in that their hydrostat is covered in thick outer skin^5^. This skin has a protective function, but elephants also utilize it to assist in gripping objects when they wrap^6^ or sweep food using the wrinkled ventral portion at the tip of the trunk^7^. Elephant skin is known for some simple gross mechanical properties, such as a cracked epidermis for thermoregulation^8^ in African elephants and entangled collagen in the dermis of the trunk for added protection and extension^9^.

The trunk’s mobility and flexibility are enabled by a highly complex musculature^10–12^, controlled by a very elaborate motor nucleus^13^. The muscles are used when an elephant reaches for objects or food^1^ and enable impressive fine motor control, enabling them to perform precise tasks such as peeling bananas^14^. They can also be controlled to manipulate air flow, allowing precise object manipulations, such as lifting a tortilla chip without breaking it using fluid suction^15^. The mechanical difficulty of using this hydrostat is obvious, as elephants can take almost a year to develop the full potential of their trunk use^16^. We see functional mechanical differences along the trunk. Specifically, the distal parts of the trunk are very dexterous and form pseudo-joints for grasping objects^7^. In contrast, proximal trunk regions play a lesser role in manipulation and are more important for support and muscular force^1^. Trunk function becomes lateralized during elephant post-natal development, and adult elephants split into left- or right-truckers according to their grasping preferences^17–19^.

African savanna elephants (*Loxodonta africana*, from here on called African elephants) and African forest elephants (*Loxodonta cyclotis*, not subject of this publication) differ from Asian elephants (*Elephas maximus*) with regards to their trunk morphology and behavior. African elephants have two finger-like protrusions on their trunk tips and tend to pinch objects with their two fingers. Asian elephants, in contrast, have only one dorsal trunk finger and tend to wrap their trunk around objects^20^.

Many mechanical differences in trunk function have been shown between African and Asian elephants and within species when comparing different developmental stages. There is little understanding, however, of the developmental factors that play a role in the functionality of the elephant trunk. Additionally, even though elephants have prominent trunk wrinkles from birth, the development of the skin of this hydrostat remains a mystery. Understanding how these wrinkles develop and change over time can help provide valuable insight into biological wrinkling and the impact of the environment and behavior on it^21^. In this study, we seek to understand the form-function ontogeny of the wrinkled trunk skin both pre-natal and through adulthood. In our analysis, we aimed to elucidate the functional and developmental characteristics of elephant trunk wrinkles. Specifically, we ask: (1) What is the number and distribution of wrinkles over the trunk in adult elephants, calves/fetuses, and across elephant species? (2) Are the elephant’s trunk wrinkles affected by trunk use and lateralization? (3) Do the skin layers differ along a single wrinkle in the trunk? (4) How do elephant trunks and trunk wrinkles develop?

## Materials and Methods

### Elephant specimens

All post-mortem specimens used in this study came from zoo elephants and were collected by the IZW (Leibniz Institute for Zoo and Wildlife Research, Berlin) over the last three decades in agreement with CITES (Convention on International Trade in Endangered Species of Wild Fauna and Flora) regulations. Specimen reports and CITES documentation for all animals included are held at the IZW. All of these elephants had died of natural causes or were euthanized by experienced zoo veterinarians for humanitarian reasons, because of insurmountable health complications. Most of the trunks used were either fixed in 4% formaldehyde solution or frozen at −20 °C. Table 1 gives an overview of the post-mortem specimens of Asian elephants (*Elephas maximus*) and African elephants (*Loxodonta africana*), along with their age.

In addition to the post-mortem samples, photographs of living elephants in zoos were also analyzed. Table 2 gives an overview of these elephants. Photographs were either taken by one of the authors at the Berlin Zoo, and the Zoo Schönbrunn, Vienna, or provided by zoo employees, collaborators, or photographers.

### Photography of trunks

Post-mortem specimens at the lab and elephants at the zoos were photographed using a Sony α 7R III camera or a Sony α 7S II with a Sony FE 16-35mm F2.8 GM E-Mount objective, a Sony FE 90 Mm/2.8 Macro G OSS objective, or a Sony FE 4.5-5.6/100-400 GM OSS zoom objective. Cameras were used handheld or mounted on a Hama “Star 62” tripod or a Manfrotto “MT190CXPRO4” carbon tripod.

### Elephant wrinkle measurements

The trunks were divided into zones: base, lateral shaft, dorsal shaft, ventral shaft, and tip (Figure S1). The base is determined as the most proximal part of the trunk, above the tusk pouch. The tip is considered the most distal portion of the trunk from the split into the two fingers in African or finger and cartilage in Asian elephants to the end of the fingers (Figure S1A-B). The shaft is the rest of the trunk between the distal tip and the proximal base, as described. Wrinkles were identified as either “major” or “minor” wrinkles, with the “major” being deeper, mostly regularly spaced, and transverse the whole dorsal or ventral part of the shaft (Figure 2A and B). The “minor” wrinkles are shallow wrinkles that partially cross the trunk with uneven spacing. In African elephants, the proximal trunk has been described to have deep wrinkles termed “folds”^1^, which, for our analysis, were counted as “major” wrinkles. For the lateralization (Figure 3C-E), we looked at the first 15 cm of the trunk from the tip on, and only “major” wrinkles of the shaft, not of the tip itself, were considered. Wrinkles were counted using the multi-point tool in ImageJ. Wrinkle position, including wavelength, was determined using ImageJ.

### Fetal development

To study trunk wrinkle and trunk development, we studied five fetal elephant specimens (n = 3 African elephant fetuses from the Naturkundemuseum Berlin, and n = 2 Asian elephant fetuses from our collection). We also made a major effort to collect photographs or drawings of elephant fetuses (n = 50 African, n = 12 Asian) from published work^22–48^. We assigned presumed embryonic ages, denoted as E*n* where *n* is the number of days according to the formulas for embryonic length^37^ or mass^37^ in early fetuses. In older fetuses (> E200) we used the mass-age formula developed by Craig 1984^30^.

### MicroCT scanning

All samples for microCT scanning were taken from trunks that were fixed in 4% formaldehyde for several months. To characterize wrinkles from different trunk regions, an Asian baby elephant trunk was cut in half sagittal and stained in 1% iodine solution for 33 days to enhance tissue contrast. The half trunk was then stained for 84 days in a lower concentration of Iodine solution. For the African baby elephant trunk, the sample was first put for 30 days in a 1% iodine solution, 30 days in a 2% iodine solution, and finally 30 days in a 3% iodine solution.

All iodine solutions were prepared by diluting 5% Lugol’s iodine in distilled water. The scans for the Asian baby elephant trunk were performed using the YXLON FF20 CT scanner (YXLON International GmbH, Hamburg, Germany) at the Humboldt University of Berlin. The African baby elephant trunk was scanned at the Museum für Naturkunde Berlin with a YXLON FF 85 CT (YXLON International GmbH, Hamburg, Germany).

### MicroCT and histology determining trunk wrinkle and amplitude and skin thickness

The amplitudes of the trunk wrinkles were taken as a trough-to-peak measurement as in a sinusoidal wave. This is a simple estimation to differentiate the wrinkled pattern in the post-mortem baby specimens. The amplitude was calculated using side views of transversely dissected trunks allowing peak-to-peak calculation of various segments along the trunk’s surface.

The wavelength of the trunk wrinkles was taken as the distance between any two wrinkles. The wrinkles were sketched as tangential lines. To calculate the wavelength and the wrinkle number, perpendicular lines were drawn from the left side to the right side of the trunk. The number of intersections was described as a wrinkle number for that segment, and the distance between each intersection is the wavelength between those wrinkles. We analyzed all the zones previously described for the average wavelength between wrinkles and the wrinkle number along the trunk.

To compare the African and Asian elephant microCTs (Figure 4A & Figure 4E) we normalized the positional information using the total trunk length. Therefore, the trunk wrinkle amplitude & wavelength are plotted on the same axis of African and Asian elephants by dividing the position along the trunk by total length, giving a dimensionless length. This means the trunk position is unitless, and a trunk position of 0 is at the proximal base of the trunk, and near the distal tip, it would have a value of 1. To perform statistical comparisons along the normalized length of the African and Asian elephants, we performed zone-wise comparisons between three primary zones: the proximal, mid-section, and distal sections. Each section has a normalized length of 0.3 of the normalized trunk length, with the proximal section ranging from 0-0.3, the midsection 0.3-0.6, and the distal section 0.6-0.9. We did not analyze the normalized trunk length of 0.9-1.0 as this portion of the microCTs did not have enough resolution to get amplitude or wavelength measurements. The three sections were averaged for each specimen and compared.

We also show histological sections from an Asian baby elephant’s trunk tip finger (Figure S2A; originally done for Deiringer et. al 2023^49^ but not shown there) to determine the histological differences between the two primary skin layers (Figure S2B-C). Samples were stained using a standard Hematoxylin-eosin stain for elephant tissue^49^ and imaged using an Olympus BX51 Microscope (Olympus, Japan) with an MBFCX9000 camera (MBF Bioscience, Williston, USA).

### Statistical analysis of wrinkle numbers

Overall, we used several different statistical tests throughout the analysis of the manuscript. To compare the total wrinkle numbers of adult Africans to adult Asian elephants, we used a Mann-Whitney U test. To compare major trunk wrinkles in adult elephants between species, we used two-sample t-tests, specifically Welch’s t-test, to compare all adults and the Pooled variance to compare females. We examined differences between dorsal and ventral portions of the trunk with a two-tailed paired t-test. To determine differences in major wrinkles on the trunk shaft, we used a two-sample t-test (Pooled variance). When determining whether the trunk tip side with the longer or the one with the shorter whiskers has more wrinkles, a Wilcoxon Signed-Rank test was performed. A second Wilcoxon Signed-Rank test revealed that wrinkle numbers do not differ overall between the left or right trunk side in African and Asian elephants. We performed a one-way ANOVA for each zone to test amplitude and wavelength differences. All t-tests were two-sided, and the null hypothesis was rejected when p < 0.05. The methods and results section clarifies each statistical test’s sample size.

## Results

We studied wrinkles on the trunk of Asian (*Elephas maximus*) and African (*Loxodonta africana*) elephants (Figure 1). We analyzed photographs of live elephants from Zoos (Table S2) and post-mortem samples that were collected in a decade-long effort by the IZW (Leibniz Institute for Zoo and Wildlife Research, Berlin) (Table S1). We looked at skin structure in relation to wrinkles using post-mortem specimens and MicroCT scans. To examine the early development of wrinkles, we studied post-mortem material from fetuses and newborns.

**Figure 1:**
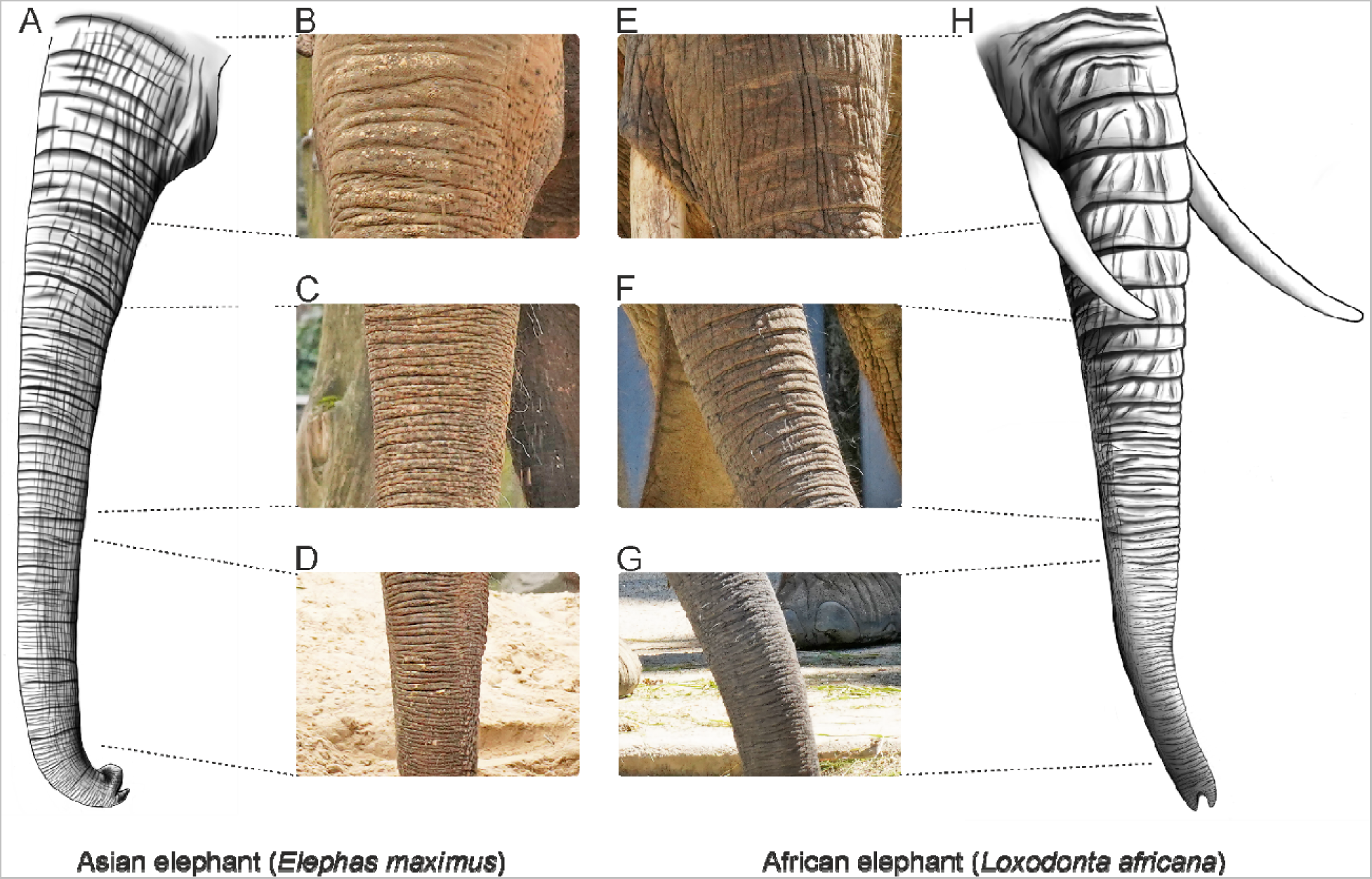
Asian (*Elephas maximus*) and African (*Loxodonta africana*) elephants differ in trunk morphology, including trunk wrinkles. **A**, Drawing of the trunk of an adult Asian elephant. Note the larger number of transversal wrinkles in the Asian elephant compared to the African elephant trunk in **H**. **B**, Proximal trunk base wrinkles of an Asian elephant. **C**, Same as **B** but for the middle part of the trunk. **D**, Same as **B** but for the distal part of the trunk. **E-G**, Same as **B-D** but for an African elephant **H**, Drawing of the trunk of an adult African elephant. Note the major wrinkles on the proximal half of the trunk that fold over each other and how they transition to the tightly packed distal trunk wrinkles. Illustrations (A, H): Cindy Ritter. Photo credit (B-D): Lena Kaufmann, Humboldt Universität zu Berlin; Zoologischer Garten Berlin, Berlin, Germany. Photo credit (E-G): Lena Kaufmann, Humboldt Universität zu Berlin; Zoo Schönbrunn, Vienna, Austria.

### Asian and African elephants differ in trunk wrinkles and overall morphology

Asian (Figure 1A) and African (Figure 1H) elephant trunks differ in their morphology, with one of the obvious differences being in the form and arrangement of wrinkles that can be found on the whole trunk from the base until the very tip. The coloration of the trunk and skin texture differs between species, with Asian elephant trunk skin looking lighter, having pinkish pigmentation, and smoother skin (Figure 1B-D). African elephant trunk skin appears greyer and more cracked (Figure 1E-G). In both species transversal trunk wrinkles are more clear and deeper than longitudinal trunk wrinkles and there are few or no oblique (non-transversal or longitudinal) wrinkles (Figure 1). We found that the number of total trunk wrinkles (major + minor) in adult Asian elephants (n = 7, out of these 5 females and 2 males; x = 155, SD = 26) is larger than in African elephants (n = 7, all females; x = 109, SD = 14; Mann Whitney U test, p = 0.007, Z = 2.69). In both species, we can see the distance between wrinkles (wavelength) decreasing towards the distal end of the trunk (Figure 1). In Asian elephants, wrinkles of the proximal trunk appear shallower than in African elephants and more irregularly spaced (Figure 1B and E, note also the schematic skin cross-section diagrams showing wrinkles in profile below the photographs). Wrinkles of the medial (Figure 1C and F) and distal shaft (Figure 1D and G) appear more densely packed in Asian than in African elephants. For a visualization of the partition of the trunk in base, shaft, and tip zones see Figure S1A. In both elephant species studied we also noticed numerous partial wrinkles both on the proximal base and distal tip of the trunks. These partial, “broken” wrinkles wrap around half of the trunk from one lateral side and after a gap often continuing shifted a little bit proximally or distally. The remainder of our paper will focus on transversal wrinkles on the dorsal or ventral trunk. We conclude that Asian and African elephants have visually distinct patterns of trunk wrinkles.

### Trunk major and minor wrinkles differ in counts and distribution between Asian and African elephants

Wrinkles were traced on photographs of elephant trunks and color-coded according to wrinkle type, which could be major or minor wrinkles (Figures 2A and B, note the schematics depicting major/minor wrinkles in both species). Major wrinkles are deeper and more regularly spaced than minor wrinkles. For details see methods and Figure 4, where microCT scans very clearly show the difference between the deep major wrinkles and the shallower minor wrinkles, often lying in between two major ones.

**Figure 2:**
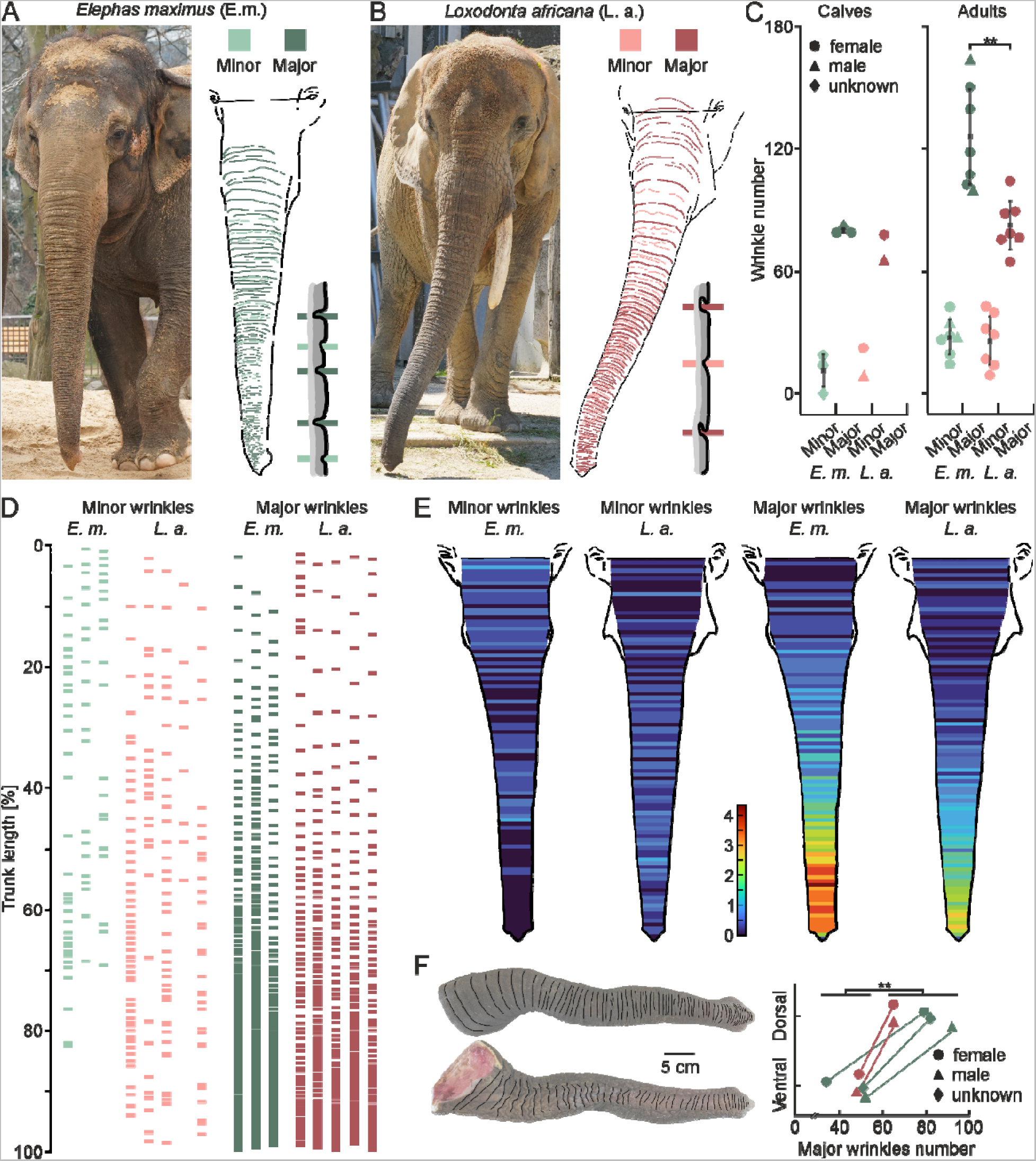
Transversal trunk major and minor wrinkles differ in counts and distribution between Asian and African elephants. **A**, Female Asian elephant Carla (Zoo Berlin) next to a tracing of minor and major wrinkles with a schematic of what minor/major wrinkles look like in Asian elephants. Major wrinkles in contrast to minor wrinkles are deeper, more regularly occurring, and regularly spaced, and transverse the whole dorsal or ventral trunk. Note the zero-line between the eyes, wrinkles proximal to this were not included in the analysis. **B**, Same as A but for female African elephant Drumbo (at the time of photograph Zoo Schönbrunn). **C**, Comparison of trunk wrinkle numbers between Asian and African elephants in calves (left) and adults (right). Asian elephant calves (n = 3) and African elephant calves (n = 2) have similar numbers of minor and major wrinkles. Asian adult elephants (n = 7) have more major trunk wrinkles (x = 126, SD = 25) than African adult elephants (n = 7, x = 83, SD = 13; two-sample t-test (12) = 4, p = 0.003, d = 2.16). Circle = female, triangle = male, diamond = unknown. **D**, Positions of minor (left) and major (right) wrinkles on trunks normalized to total trunk length in individual female Asian (n = 3) and African (n = 5) elephants. **E**, Heatmaps showing the distribution of minor (left) and major (right) trunk wrinkles in Asian (n = 3) and African (n = 5) elephants, based on **D**. Asian elephants have on average more minor wrinkles in the proximal part of the trunk whereas in African elephants minor wrinkles are more spread over the rest of the trunk. Asian elephants have on average more major wrinkles in the distal half of the trunk with a particularly high density in the region where they bend when wrapping objects. Blue shows a low average density of wrinkles and red a high average density of wrinkles at this position of the trunk. **F**, On the left a photograph of an African elephant calf trunk showing tracings of major wrinkles on the dorsal (upper) and ventral (lower) sides of the trunk. On the right a comparison of major wrinkles on the dorsal and ventral sides of the same trunk in Asian (n = 3) and African (n = 2) elephants. There are significantly more wrinkles on the dorsal (M = 77, SD = 12) than on the ventral sides of the trunks (M = 47, SD = 7; two-tails paired t-test(4) = 5.1, p = 0.007, d = 2.27). Circle = female, triangle = male, diamond = unknown. The line between symbols represents both symbols being part of the same trunk, one being the winkle number on the dorsal and one the wrinkle number on the ventral side. Photo credit (A): Lena Kaufmann, Humboldt Universität zu Berlin; Zoologischer Garten Berlin, Berlin, Germany. Photo credit (B): Lena Kaufmann, Humboldt Universität zu Berlin; Zoo Schönbrunn, Vienna, Austria. Photo credit (F): Lena Kaufmann, Humboldt Universität zu Berlin.

In elephant calves, we did not find differences in major or minor wrinkle numbers (Figure 2C). Adult Asian elephants (n = 7), however, have significantly more dorsal major trunk wrinkles (x = 126.43, SD = 25.3) than African elephants (n = 7, x = 83, SD = 12.87; two-sample t-test (12) = 4.05, p = 0.003, d = 2.16) but similar numbers of minor trunk wrinkles (Figure 2C). When controlling for sex and comparing only female elephants, the difference in major wrinkle numbers between Asian (n = 5, x = 124.2, SD = 20.66) and African elephants is even stronger (two-sample t-test (10) = 4.28, p = 0.002, d = 2.51). In Asian elephants, wrinkle numbers increase from on average 91 (SD = 8.19) total wrinkles in the calves (n = 3) to ~155 (SD = 26.25) total wrinkles in the adults (n = 7). This change is mostly due to the increase in major wrinkles from 80 (SD = 1.73) in the calves to ~126 (SD = 25.3) in the adult Asian elephants. For the African elephants, on the other hand, the change in total wrinkle number from on average ~87 (SD = 19.09) in the calves (n = 2) to ~109 (SD = 14.28) in the adults is a bit smaller. Major wrinkles increase in African elephants from ~72 (SD = 9.19) in the calves to on average 83 (SD = 12.87) in the adults (Figure 2C).

The minor wrinkles are denser in the Asian elephant’s proximal part of the trunk, whereas in African elephants they are more spread out than in the Asian and denser in the distal part of the trunk (Figure 2D and E). In both species, the density of major wrinkles increases towards the trunk tip, with the average density of major wrinkles at the distal third of the trunk being even higher in Asian elephants than in African elephants (Figures 2D and E). In both species (n = 5) we found a significantly greater number of major wrinkles dorsally (x = 76.6, SD = 10.4) than ventrally (x = 46.8, SD = 6.55; paired t-test (4) = −5.07, p = 0.007, d = 2.27; Figure 2F) in both Asian (n = 3, all <5 y. o.) and African (n = 2, one adult and one calf).

A more in-depth analysis of wrinkle numbers separated in different trunk zones (base, shaft, or tip; see Figure S1A and B) revealed a difference in major wrinkles on the trunk shaft, with adult Asian elephants (n = 7) having a statistically greater number of major trunk shaft wrinkles (x = 115.29, SD = 26.2) than adult African elephants (n = 7, x = 70.43, SD = 12.55; two-sample t-test (12) = 4.09, p = 0.002, d = 2.18; Figure S1C). No differences between species were found in the number of wrinkles on the trunk base or tip, independently of pooling or not pooling major and minor wrinkles or female and male elephants. There is also no significant difference between the two species in minor trunk wrinkle numbers of the trunk shaft. In calves of both species, we found comparable numbers of major and minor wrinkles, so it is notable that Asian elephants gain more wrinkles during their life than African elephants (Figure S1C). The differences in wrinkle numbers between the two species reflect differences in the number of major wrinkles, but not minor wrinkles (Figure 2C and Figure S1C).

### Trunk wrinkle number is lateralized

Trunk wrinkle numbers continue to increase throughout ontogenetic development. We also discovered that through mechanical usage elephants develop wrinkles on their lateral trunk sides (Figure 3A-B). Almost all adult elephants show marked left-right asymmetries in whisker length as shown by Deiringer et al. (2023)^49^. Elephants use the distal third of their trunk to wrap food or other objects and the whiskers are longer on the side they are wrapping towards, designating the “trunkedness” (Figure 3C). This whisker asymmetry is age- and use-dependent as elephant calves are born without it and will develop a favored side along with their trunk control. Whisker abrasion will appear on the opposite side of the one wrapped towards, as it will be more often in contact with the ground. We identified the “trunkedness” of all our specimens, as illustrated with the trunk tip of an African elephant in Figure 3C. Whiskers were longer on its right trunk side; thus, it presumably was a right-trunker, preferentially wrapping towards its right side.

**Figure 3.**
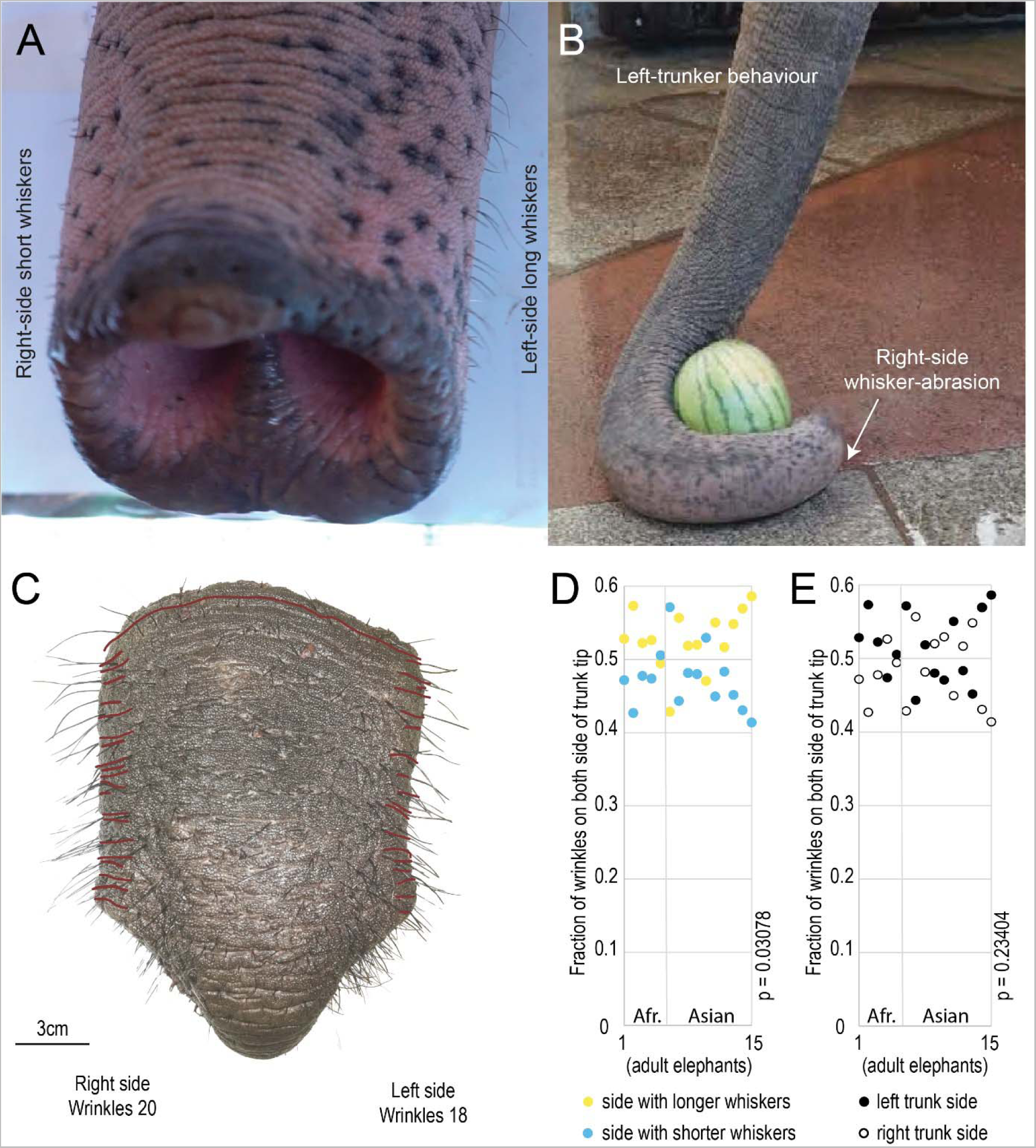
Left- and right-trunkers have more wrinkles on the left and right trunk sides, respectively. **A**, Image of a left-trunker with the trunk tip and distal trunk shaft of an Asian elephant. Note the shorter whiskers on the right side and the longer whiskers on the left side of the distal trunk shaft. Modified from Deiringer et. al^49^. **B,** Image showing a left-trunker behavior of an Asian elephant reaching for a watermelon. The image displays a trunk wrapping to the left with the right-side of the distal shaft of the trunk in contact with the ground which results in whisker abrasion. Modified from Deiringer et. al (2023)^49^. **C**, African Elephant trunk tip and distal trunk shaft, in red the major wrinkles on the trunk shaft that were counted. The red line crossing the tip is the last wrinkle counted. **D**, Univariate plot of the fraction of wrinkles (normalized to the total count on both trunk sides) on the trunk side with shorter or longer whiskers. Trunk function is lateralized in elephants and so-called left-trunkers, who preferentially grasp towards the left side, have shorter whiskers of the right side of the distal trunk shaft^49^; the reverse is true for right-trunkers. We observed ~10% more wrinkles on the longer whisker side. In yellow the wrinkles fraction on the longer whisker side and in blue the wrinkles on the shorter whisker side. One dot is one animal, yellow and blue for the same animal plotted on the same axis. The wrinkles counts are normalized by the total wrinkles count. (Wilcoxon test, z-value = −2.1583, w-value = 22, p = 0.03078, d = −0.55727) (5 adult African elephants, 9 adult Asian elephants). **E**, Univariate plot of the fraction of wrinkles (normalized to the total count on both trunk sides) on the left or right side of the trunk. The full dots are the wrinkles fraction on the left side and the empty dots are the wrinkles on the right side. One dot is one animal, for the same animal the dots are plotted on the same axis. The wrinkles counts are normalized by the total wrinkles count. (Wilcoxon test, z-value = −1.1927, w-value = 39, p = 0.23404, d = −0.30795) (5 adult African elephants, 9 adult Asian elephants). Photo credit (A, B, C): Lena Kaufmann, Humboldt Universität zu Berlin.

In both, Asian (n = 10) and African (n = 5) elephants, we observed ~10% more wrinkles on the distal trunk shaft side with the longer whiskers. This bias of having more wrinkles on the trunk side with longer whiskers (x = 0.52, SD = 0.04) than on the shorter whiskers side (x = 0.48, SD = 0.04) was systematic and significant (Wilcoxon Signed-Rank, p = 0.03, z-value = −2.16, w-value = 22, d = −0.56; Figure 3D). In contrast, we did not observe a systematic difference in trunk side wrinkle numbers as a function of left (x = 0.52, SD = 0.05) vs. right trunk side (x = 0.49, SD = 0.05; Wilcoxon Signed-Rank test, p = 0.23, z-value = −1.19, w-value = 39, d = −0.31), nor with a species difference (Figure 3E). Taken together, this indicates that differences in trunk side wrinkle numbers are due to the individual’s “trunkedness”, being a “left-“ or “right-trunker”, or, in other words, behavioral preferences shape the morphology or the trunk.

### Skin layers shift along the wrinkled skin of elephant trunks

To determine the amplitudes of wrinkles along the trunk, we analyzed preserved trunk specimens. We performed microCT scans of iodine-stained trunks of Asian and African elephant calves to visualize trunk wrinkles and underlying skin structure. These scans provided high-resolution images of entire elephant trunks (Figure 4). Major and minor wrinkles were readily visible in volume renderings (Figure 4A) and parasagittal sections (Figure 4B) of an Asian baby elephant trunk. As we noted before in adult elephant trunks, wrinkle frequency increased from proximal to distal (Figure 4C and D).

**Figure 4.**
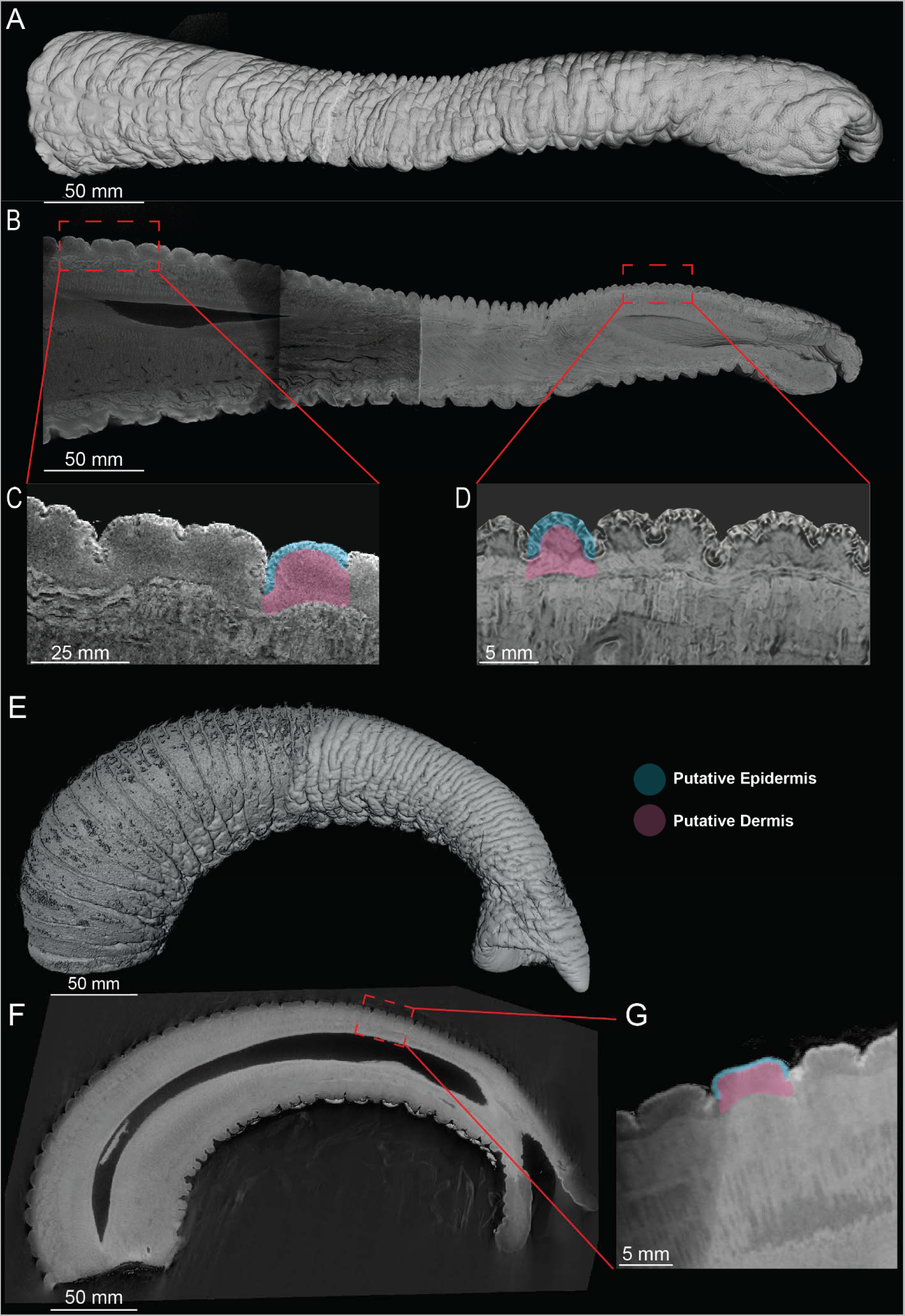
Visualization of trunk wrinkles in microCT scans of an Asian and an African baby elephant trunk. **A**, Volume rendering of a µCT scanned Asian baby elephant trunk. **B**, Sagittal slice of an Asian baby elephant trunk. **C**, High magnification view of proximal dorsal wrinkles in the Asian baby elephant trunk. Note the constant thickness of the putative epidermis, while the putative dermis is getting thinner in the throughs of the wrinkles. The two primary load-bearing layers of skin are highlighted the epidermis in blue, and the dermis in pink. **D**, High magnification view of distal dorsal wrinkles in the Asian baby elephant trunk with highlights of the two primary load-bearing skin layers. Note the constant thickness of the putative epidermis, while the putative dermis is getting thinner in the throughs of the wrinkles. **E**, Volume rendering of a µCT scanned African baby elephant trunk. **F**, Sagittal slice of an African baby elephant trunk. **G**, High magnification view of dorsal wrinkles in the African baby elephant trunk. Note the constant thickness of the putative epidermis, while the putative dermis is getting thinner in the throughs of the wrinkles. The two primary load-bearing layers of skin are highlighted similarly to **C** and **D**.

It appears that the morphology of the different skin layers differs along the length of a wrinkle. In the Asian baby elephant, the outer layer of the skin, the epidermis (blue highlight), appears to be constant throughout a wrinkle, however, the dermis (pink highlight) becomes thicker in between the troughs and is thinner in the trough of a wrinkle (Figure 4C-D). Thus, trunk major wrinkles are not mere creases of the skin but show clearly non-homogenous skin layers along their length. In particular, because of the reduction of parts of the inner skin in the trough, the skin is quite thin and presumably also more flexible. Minor wrinkles appeared to be slight indentions in the epidermis, like folding of the major wrinkle crest on itself. Very similar observations were made on volume renderings (Figure 4E) and parasagittal sections (Figure 4F) of an African baby elephant trunk. Similar to the Asian baby elephant, the epidermis of the African baby elephant was observed to be of near-constant thickness throughout major wrinkles. At the same time, we found the dermis layer to change between the peak and troughs of the major wrinkles (Figure 4G). For a more detailed histological analysis of skin layers in the trunk tip see Figure S2. We conclude that the morphology of the major wrinkles in African and Asian elephants differs along the length of the wrinkle.

### Wrinkle and amplitude species differentiation in microCT

We used measurements of individual wrinkles of the microCT-scanned specimens to perform a more in-depth analysis of wrinkle wavelengths, the distances between two major wrinkle troughs, and amplitudes or depths. In both Asian and African elephants, the wavelength between dorsal trunk wrinkles was highest at the proximal portion of the trunk and decreased towards the tip (Figure 5A). In comparing the zones of the African and Asian elephants, we see African elephants have significantly larger wavelengths (Figure 5A) in the mid-section (x = 5.98, SD = 1.43, F = 32.93, p < 0.001) and distal section (x = 3.29, SD = 0.88, F = 6.98, p = 0.01) compared to Asian elephant’s mid-section (x = 3.64, SD = 0.86) and distal section (x = 2.68, SD = 0.56). This is consistent with other results, as larger wavelengths indicate fewer wrinkles along the trunk. At the distal tip, the wavelengths are nearly equivalent, with both elephants having wavelengths around 3 mm (Figure 5A). In analysis of the amplitude differences between the species we see that Asian elephants have significantly higher amplitudes along the whole trunk (Figure 5B) including proximal (x = 2.7, SD = 0.42, F = 22.5, p < 0.001), midsection (x = 1.86, SD = 0.35, F = 44.68, p < 0.001), and distal (x = 1.45, SD = 0.48, F = 11.87, p = 0.001) sections. This is compared to the African elephant with amplitudes decreasing by only by nearly 35% from the proximal section (x = 1.62, SD = 0.32) to the distal portion (x = 1.05, SD = 0.22).

**Figure 5:**
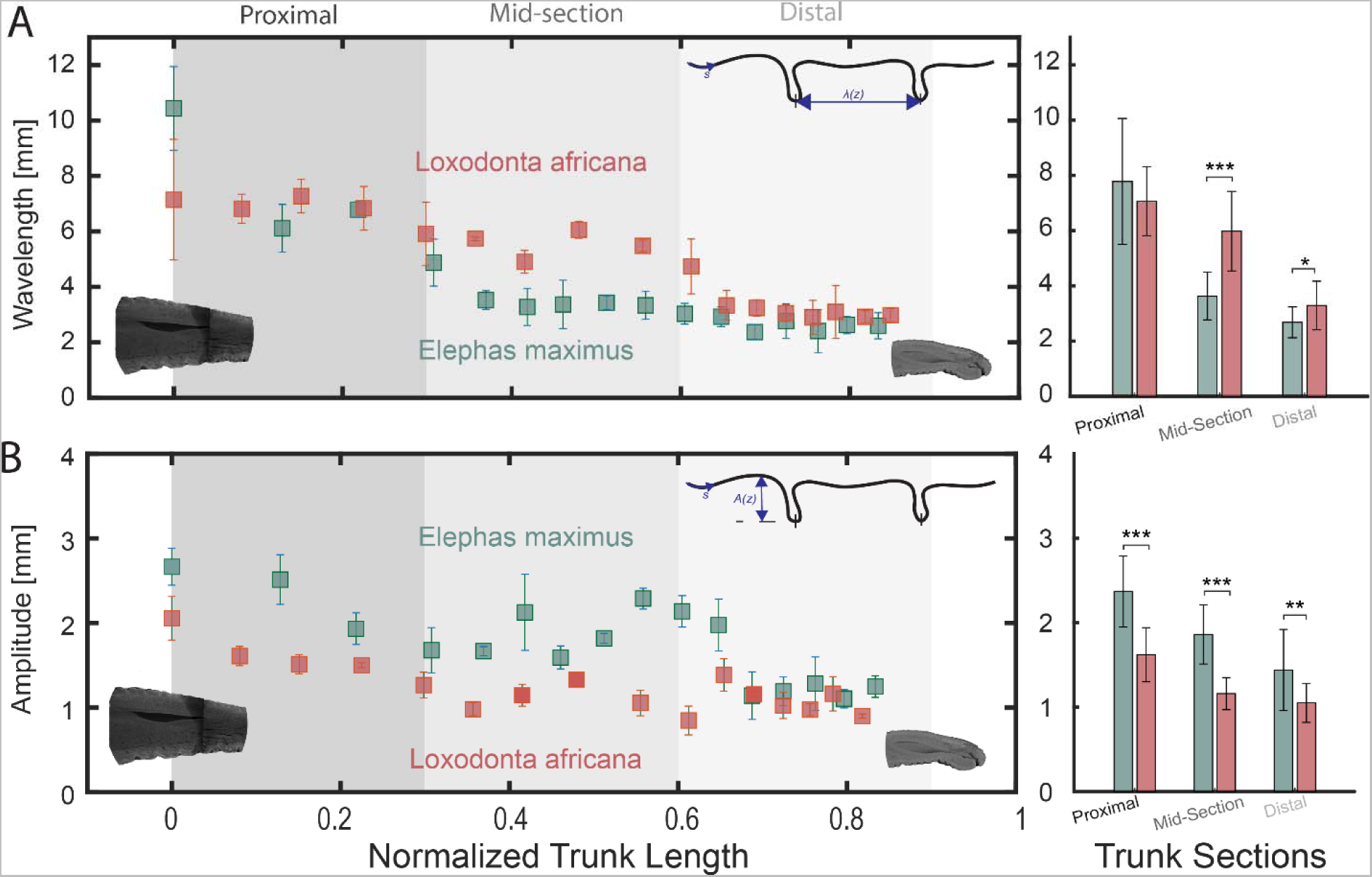
Wrinkle amplitude and wavelength of African and Asian elephant baby trunks. **A**, Wavelength of an Asian baby elephant (*Elephas maximus*; green) trunk and an African baby elephant (*Loxodonta africana*; orange) trunk, taken from the microCT-scans in Figure 4. Averages of three different proximal, mid-section, and distal sections are shown with shaded sections where the average of the wrinkle amplitude and wavelength are compared statistically in each section. Statistics for the three sections are shown on the right. **B**, Peak-to-peak amplitude of an Asian baby elephant (green) trunk and an African baby elephant (orange) trunk, taken from the microCT scans in Figure 4. The average of different zones is shown similarly to (A) with comparisons between the species. Statistics for the three sections are shown on the right.

### Fetal trunk and trunk wrinkle development

We examined how trunk wrinkles develop and how they relate to trunk development in general. To address this issue, we studied Asian (n = 2) and African elephant (n = 3) fetal specimens, as well as published photographs or drawings of elephant fetuses (n = 12 Asian, n = 50 African). We then assigned embryonic (E) ages to these specimens, as detailed in the methods section.

Figure 6A shows schematic drawings of different stages of fetal African elephant heads, their trunks (black), their trunk wrinkles (gray), and their upper (red) and lower (green) lip. The elephant trunk develops from a large nose primordium. Wrinkles are initially added rapidly and then gradually. A schematic overview is given for wrinkle development (Figure 6B) and lip development (Figure 6C). The upper lip-nose-fusion occurs rapidly between embryonic days E100 and E130.

**Figure 6:**
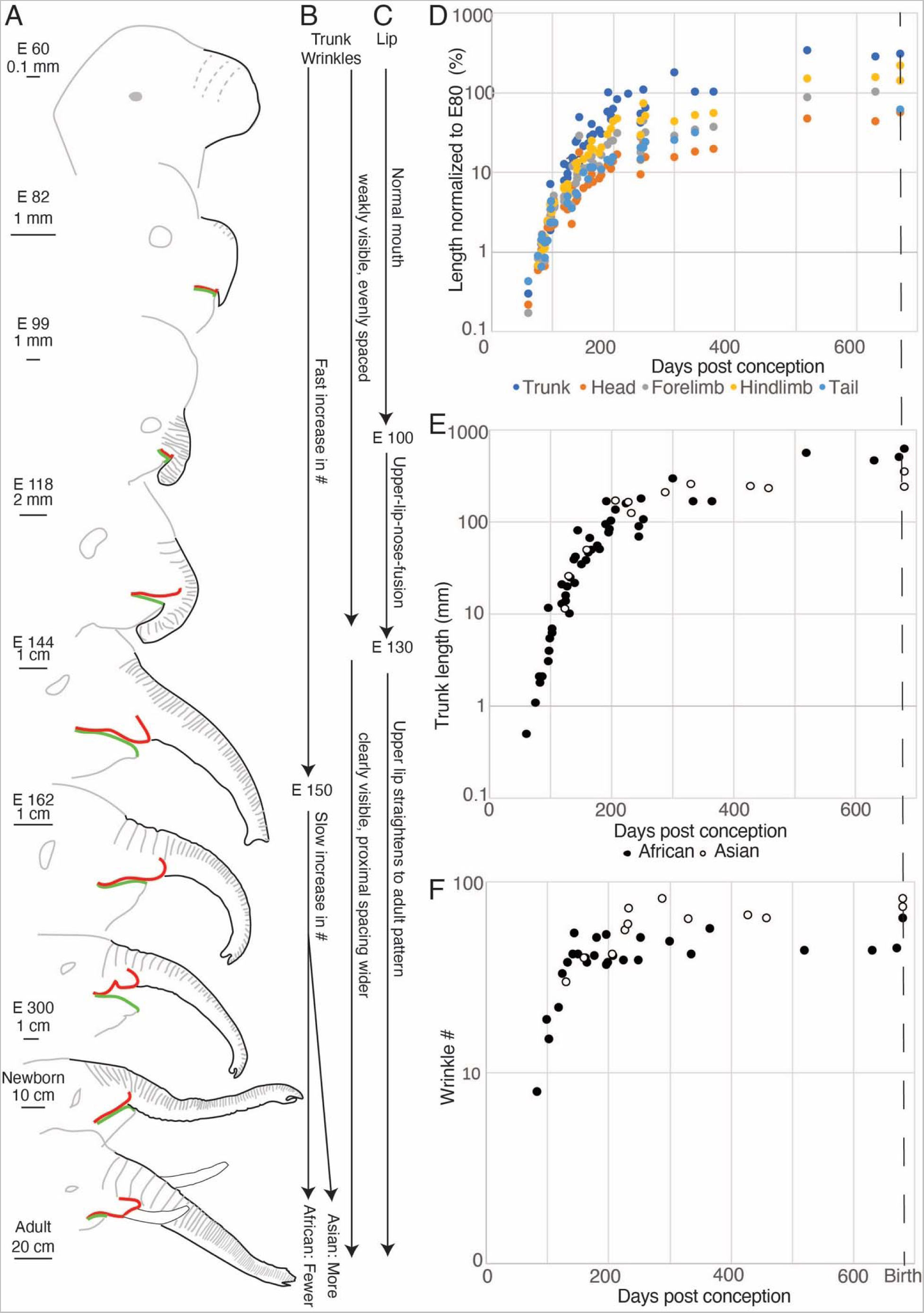
Fetal trunk and trunk wrinkle development. **A**, Schematic drawings of African elephant fetuses. Gray, outline of the head and wrinkles; gray dashed, putative wrinkles; black, outline of the trunk; green, lower lip; red, upper lip. Fetuses were redrawn from references following references^22–48^. E# indicates the embryonic age of the photo with the number indicating days. **B**, Schematic of stages of fetal wrinkle development in African and Asian elephants. **C**, Schematic of stages of fetal lip development in African elephants. **D**, Relative length growth of various body parts in African elephants. Length was normalized to the length of the respective body part in E80 fetuses and is given in %. The trunk grows more than other body parts and the accelerated growth occurs mainly between E60 and E150. **E**, Trunk length growth in African and Asian elephants is similar. **F**, Wrinkle development in African and Asian elephants. Wrinkle number increases in sharply different phases: Between E80 and E130 there is an exponential increase in wrinkle number with a doubling time of ~20 days. After E130 addition of wrinkles is slow, but slightly faster in Asian than in African elephants. Note that wrinkles could only be counted in a subset of fetuses.

The trunk shows more length growth than other elephant body parts; this faster growth occurs early (E60-E150, Figure 6D). The fetal trunk length growth pattern is similar in Asian and African elephants (Figure 6E). A log plot of wrinkle numbers against fetal age reveals that wrinkles develop in two sharply different phases (Figure 6F). Between E80 and E150 there is an exponential growth of wrinkle numbers with a doubling time of about 20 days, after that addition of wrinkles is slow, and slightly faster in Asian elephant fetuses than in African ones. The number of wrinkles on fetal elephant trunks between E200 and birth in Asian (n = 8, x = 63.63, SD = 11.82) and African elephants (n = 11, x = 44.36, SD = 5.95) is plausibly continued in the total number of wrinkles we found in Asian (n = 3, x = 91, SD = 8.19) and African (n = 2, x = 86.5, SD = 19.09) baby elephants, based on our laboratory specimen as well as photographs from zoos (Figure 2C).

Adult African elephants have two fingers at the tip of their trunk a dorsal (top) and a ventral (bottom) finger. In contrast, Asian elephants only have a dorsal finger and a ventral cartilage stump. We characterized the development of the trunk tip and the fingers in a schematic (Figure 7A). The trunk first grows as a stump. Then around embryonic day 130, the ventral finger grows out in African elephants, whereas Asians grow out a bulbous ventral trunk tip structure. The fact that the ventral finger grows out first is shown in Figure 7B for African elephants. Specifically, we observed that the ventral finger tends to be longer between E120 (before that there are no fingers) and E200. Dorsal finger development follows a slight delay in both species and differs in time course between Asian and African elephants. In African elephants, finger growth goes through a brief initial exponential length increase, after which the finger grows slower and more gradually (Figure 7C), our data were insufficient for a detailed assessment of finger growth patterns in Asian elephants.

**Figure 7:**
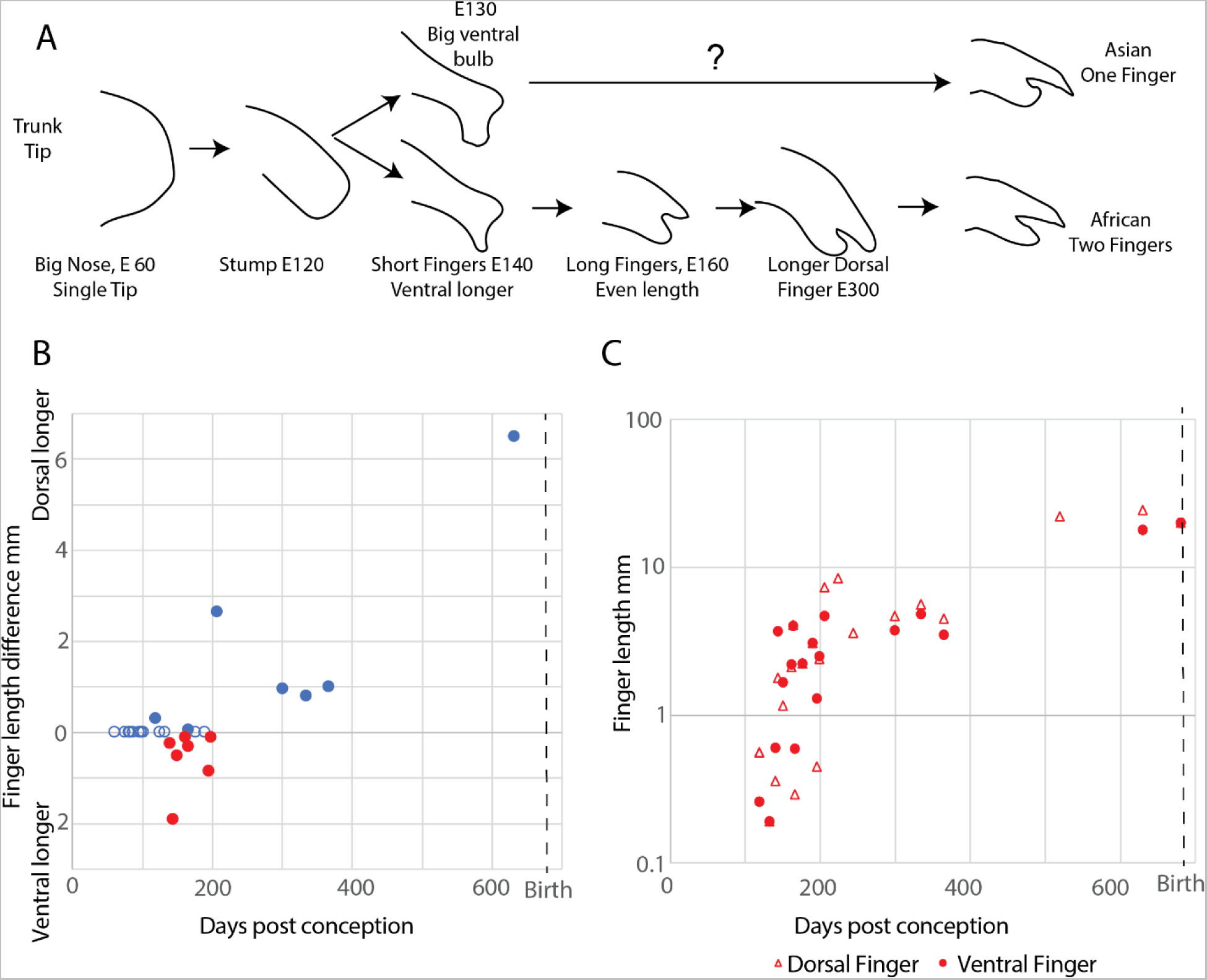
Fetal trunk finger development in Asian and African elephants. **A**, Schematic of stages of trunk tip development in African and Asian elephants. **B**, Length difference of dorsal and ventral trunk finger in African elephants throughout fetal development. Pre E120 there are no fingers, then the ventral finger is longer (highlighted as red dots) and after E200 the dorsal finger takes over. **C**, Fetal dorsal and ventral finger growth in African elephants. Finger growth goes through a brief exponential phase (E130-E180), after which finger growth slows down. The length of both the dorsal and ventral fingers could not be determined in all specimens. Note that the ventral finger (circles) tends to be longer than the dorsal finger (triangles) in early fetuses and shorter than the dorsal finger in older fetuses.

We conclude that the trunk is the fastest growing body part of elephants and that wrinkles are added in two steps, a first exponential growth step, and a second slower addition step, which differs between Asian and African elephants.

## Discussion

We assessed elephant trunk wrinkles and their pre-, as well as post-natal development by photography, microCT imaging, analysis of post-mortem specimens, and literature review in a substantial sample of fetal, newborn, and adult African and Asian elephants. We find the trunk wrinkles of African and Asian elephants to be different in several aspects.

### Differences in wrinkles are tied to genetic, behavioral, and environmental factors

Specifically, adult Asian elephants have about 1.5 times more trunk wrinkles than adult African elephants, due to an extensive addition of major wrinkles in Asian elephants during their lifetime development from calf to adult. This trend begins already at fetal stages and continues throughout postnatal development, turning into a significant difference in adult elephants. Additionally, even though we find a closer spacing of wrinkles in the distal than in the proximal trunk in both species, the density of major wrinkles on the distal third of the trunk is much higher in adult Asian than in adult African elephants.

Taken together, these results could indicate, that species differences in trunk wrinkles and trunk wrinkle morphology might have a genetic component – we already see slight differences in fetal and early postnatal stages, and differences in adults could very well also be partially genetically determined. We would also like to propose, however, that specific behavioral adaptations of Asian and African elephants contribute to the effects we see here. Asian elephants have only one finger at the tip of the trunk and a cartilage bulb on the ventral side of the tip, their preferred trunk behavior e.g. when feeding is to wrap with the distal third of the trunk^20^. This “trunk wrapping zone” in Asian elephants is also the trunk region in which we found the highest density of major wrinkles, as described above. It has been shown before that elephants form pseudo-joints with their trunks, at the very same trunk region^7^. We are suggesting that the wrinkles in the distal third of the trunk facilitate bending and wrapping and are making the formation of pseudo-joints possible. African elephants have two fingers at the tip of their trunks and prefer to pinch with their trunk tip when feeding or picking up objects within a certain range of size and form^20^. Over the course of the last years, our behavioral observations at various zoos confirmed these differences in trunk behavior.

Another factor influencing trunk wrinkling might be environmental conditions. African savanna elephants and Asian elephants are adapted to distinct environmental niches, with African elephants primarily living in dry environments^50^ and Asian elephants living in wet environments. Humidity has been shown to have an impact on human skin, with humans developing more wrinkles after transitioning from a high-humidity to a low-humidity environment^51^. The increased wrinkling in Asian elephants might be adaptive in high-humidity environments, whereas African elephants occur primarily in low-humidity environments. In investigations of thermoregulation, it has been found that Asian elephants have a higher skin temperature^52^, and higher temperatures can lead to greater wrinkling of the stratum corneum layer^53^. These thermal and environmental differences could lead to an increase in wrinkle numbers in Asian elephants.

Both, Asian and African elephants, have more major wrinkles on the dorsal side of the trunk than on the ventral side. It should be noted that most of the specimens analyzed here were <5 years old, so we think this difference is more strongly predetermined and most likely use over a lifetime is a negligible factor. This could be related to a different function of dorsal and ventral trunk wrinkles. It has been shown that the distal dorsal part of the trunk contributes the most to trunk stretching and that the ventral side stretches comparably little when the trunk is extended^1^. The distal ventral trunk has been described to be used in sweeping food together^9^ and most trunk manipulation movements are accomplished with gripping and grabbing on the ventral side^54^. Specifically, the trunk section just before the trunk tip is used in holding food or other objects, often between the lateral skin ridges that go along the ventral trunk and that have a very high density of whiskers in this distinct trunk part^49^. We are suggesting that the dorsoventral difference in trunk wrinkles can be explained by the dorsal wrinkles contributing strongly to the trunk’s ability to stretch, while the ventral wrinkles are especially important for improved grip.

### Trunk lateralization drives wrinkle differences

When looking at the most distal 15 cm of the trunks of adult Asian and African elephants, we found a difference of 10% in wrinkle numbers between the left/right side of the trunk shaft correlating with the individual’s “trunkedness”. Both Asian^18^ and African^19^ elephants exhibit lateralization, or “handedness”/”trunkedness”, with their trunks, meaning they will prefer a direction when executing complex motion tasks. This lateralization means the trunk is contacting the ground more often with one side, causing additional force and abrasions, e.g. of the whiskers, at this side of the distal trunk. Additionally, lateralization in elephants indicates that they curve the trunk to wrap and pick up objects to a specific side, left or right, making them “left-“ or “right-trunkers”^7^. Our results show that in adult elephants there are more wrinkles on the side of the distal trunk the elephant is preferentially bending or wrapping the trunk towards. To give an example, a “left-trunker” would preferably wrap their trunk towards the left of their body and perform left-oriented behaviors with the trunk, thereby frequently compressing the left side of their distal trunk and stretching the right side at the same time. The fact that there are more wrinkles on the trunk side towards which elephants preferentially wrap and that is, therefore, more often compressed points to the increase in wrinkles being a result of long-term lateralized use of the trunk.

The differences in wrinkle numbers between sides of the distal trunk are independent of species and there is no overall difference between the left and right side when we look at our samples taken together. The absence of such a left/right difference is in line with the fact that in elephants, in contrast to humans, there is no overall population-wide bias toward one side in trunk lateralization^11^. We had one case of an ambidextrous African elephant with no whisker or wrinkle difference, reinforcing our theory of the modification of the wrinkle pattern based on a user-dependent experience. If the elephant does not favor its left or right trunk side, there won’t be an abrasion of whiskers nor a more frequent compression on its trunk skin on one side. Without this frequent compression, the trunk would not get more wrinkles. In the elephant calves that we looked at, the numbers of trunk wrinkles were comparable for both sides of the trunk. Because trunk lateralization emerges with the functional ability of the trunk^55^, and it takes nearly two years to gain full control of the trunk^56^, our data indicate that trunk wrinkle patterns are affected by use. Elephants use their trunks daily to grab objects and eat nearly 200 kg of food^57^, potentially leading to around a million compression cycles a year due to lateralization. It has been shown that compression of film-substrate systems leads to a mismatching of the modulus, similar to the skin modulus, and creases and wrinkles form^58^. Therefore, millions of cycles of lateralized compression could easily lead to an increase in wrinkles.

### Changes in skin layer thickness might contribute to the functionality of wrinkles

In both elephant species, we see that in the troughs of the wrinkles, the total skin thickness decreases by coarsely a factor of two. Previously it has been described that the troughs of the wrinkles on African elephant trunks are the primary stress concentration zone when stretching^1^. In the MicroCT scans of the Asian and the African elephant calves we can see that trunk skin layers shift, with the dermis shrinking in the troughs of the wrinkles.

Wrinkle formation is often described as an instability that develops from stress, displacement, or bending that acts on a non-wrinkled surface^59^. In humans, wrinkling is related to the skin elasticity and weakening of the upper dermis^60^. In the case of elephants, it appears they have wrinkles that form from instabilities like lateralization, but as we show, they also develop wrinkles before birth. Therefore, in these elephant trunks, instead of a mechanical instability forming these wrinkles before birth, another explanation is morphological instabilities from skin layer differences along the length of a wrinkle. Having surface morphological instabilities in materials produces different types of wrinkling behavior in bilayer tubes^61^ it looks as if shifts in skin layer thickness in elephant trunks help with the formation and function of wrinkling. We also see partial wrinkles, which we denote as “broken wrinkles” that are mostly spanning from the lateral trunk towards the middle of the dorsal trunk shaft where they either just have a gap or a gap and a shift up or down. They do not continue to the ventral portions. These broken wrinkles may operate to allow additional flexibility in lateral manipulation events, assisting with enlarging the surface area of interaction with objects, similar to how specialized wrinkles in the intestine provide an enlarged surface through a broken wrinkled type of mechanism^62^.

We found in our analysis that the African baby elephant trunk has larger wavelengths than the Asian baby elephant trunk in the medial towards distal part (Figure 5A); this is consistent with the rest of our findings, as larger wavelengths along the trunk indicate fewer wrinkles. We also found significant differences in wrinkle amplitude between the two specimens, with the Asian baby elephant trunk having deeper wrinkles than the African baby elephant trunk (Figure 5B). The amplitude differences are already seen around birth, so they seem to mark a predetermined difference between the two species, and they could impact the ability to stretch out to reach far objects.

### Elephant trunk wrinkles develop early in fetal ontogeny

We analyzed elephant trunk development and trunk wrinkle development in fetal specimens and photographs of elephant fetuses. We find that the trunk shows exceptional growth in early pregnancy (E60-E150) that exceeds that of other body parts. These findings align with earlier conclusions on trunk development from transrectal ultrasound imaging^37^. Even the earliest elephant (~E60) fetuses, for which photographs are available have a large nose with periodic irregularities, possible precursors of wrinkles. Our analysis indicates a bipartite developmental pattern of trunk wrinkles. First, we observe an exponential increase in wrinkle number with a doubling of wrinkle number approximately every 20 days; this period coincides with the time of very fast trunk extension. Then, at ~E130 the increase in wrinkle number slows down sharply and continues to be slow into adulthood.

Our observations also provide a staging/dating of the upper lip nose fusion^33,34^, which appears to occur between postnatal E100 and E130, i.e. roughly in the fourth month of pregnancy. Finally, we find that trunk tip development trails trunk extension and that the trunk first grows out as a stump. Then, after E120 the fingers develop, whereby the ventral finger extends first. The bulbous ventral trunk tip of Asian elephants is present at an early point (~E130).

## Conclusion

Wrinkles and creases improve the ability of soft biological materials to bend^63^, which might explain many of our findings, in particular the proximal-distal, dorsoventral, lateralized, and species differences in wrinkle distribution. The high density of wrinkles in the Asian elephants’ ‘trunk wrapping zone’ shows how trunk wrinkles and their unique form in African and Asian elephants could contribute to the phenomenally flexible actuation of trunks. Our analysis extends earlier work on the wrinkle structure of elephant skin^1^ and gives insights into the development of the largest extant land mammals.

## Data Access Statement

The data from the paper is included in a Dryad repository for specific wrinkle numbers for each animal. Additional information for the reviewers can be found in the supplemental information which contains detailed tables of wrinkled number. The reviewer data set for the Dryad repository is included here: https://datadryad.org/stash/share/jvARnmRLbIdMtew1FskdF6XCUGEldMr2G4Jt5p6tH7I.

## Ethics Statement

All material and morphology measurements were taken from photos of previously dissected species. Specific information about each of the individuals analyzed is included in the supplement.

## Acronyms

E. m.: Elephas maximus
L. a.: Loxodonta africana
E#: Foetal age where # is how many days post presumed conception

## Acknowledgments

We thank the Berlin Zoological Garden and in particular Rolf Becker, Rouven Schulze, Konstantin Becker, and Lucas Baum. Petra Prager with her extensive collection of photographs of elephants in zoos, as well as several zoological institutions contributed. In particular the Berlin Zoo (Germany) and the Zoo Schönbrunn (Vienna, Austria), as well as Zoo Augsburg (Germany), Opel-Zoo Kronberg (Germany), Zoo Poznan (Poland), Tierpark Hagenbeck (Germany), the Elefantenhof Platschow (Germany), and the Tbilisi Zoo (Georgia). We also thank our collaborators Ani Shubitidze, Lennart Eigen, Undine Schneeweiss, Luke Longren, and Eduard Maier for their precious help in different parts of this project. AKS thanks the Alexander von Humboldt Foundation, Max Planck Society, and the International Max Planck Research School for Intelligent Systems for the Support.

## Author Contributions

A.S., L.K. and N.R. contributed equally.

Conceptualization: A.S., N.R., L.K. M.B. Data Curation: N.R., L.K., M.B., T.H. Formal Analysis: A.S., N.R. L.K. Funding acquisition: M.B. Investigation: A.S., N.R., L.K. Methodology: A.S., N.R., L.K., M.B. Project Administration: T.H., M.B. Resources: C.R., T.H., M.B. Software: A.S., N.R., L.K. Supervision: T.H., M.B. Validation: A.S., N.R., L.K., M.B. Visualization: A.S., N.R., L.K., C.R. Writing – Original Draft: A.S., N.R., L.K., M.B. Writing – Review & Editing: A.S., N.R., L.K.

## Supplement

**Figure S1:**
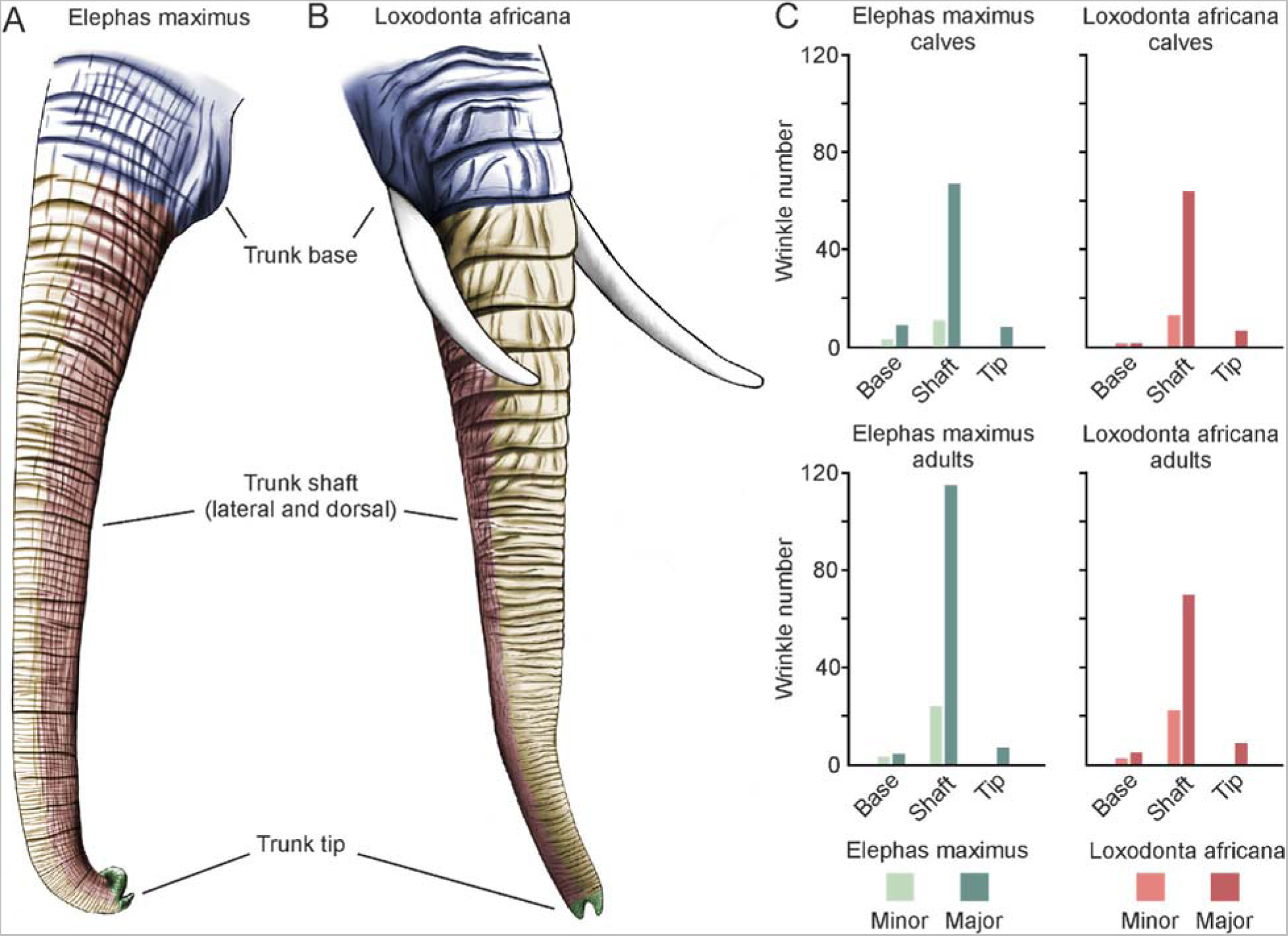
Differences in trunk wrinkle numbers between Asian and African elephants according to trunk zones and as a result of postnatal ontogeny. **A**, Drawing of an Asian elephant (*Elephas maximus*) trunk, separated in three distinct zones, base, shaft (lateral and dorsal), and tip of the trunk. Drawing taken from Figure 1. **B**, Same as in **A** but for an African elephant (*Loxodonta africana*) trunk. **C**, Minor and major wrinkle numbers in Asian (left) and African (right) elephant calves (top) and adults (bottom). Asian elephant calves (base and tip n = 3, shaft n = 4) and African elephant calves (all trunk zones n = 2) have similar numbers of both minor and major wrinkles. Adult Asian elephants (base and shaft n = 7, tip n = 10) and adult African elephants (base and shaft n = 7, tip n = 9) have comparable numbers of minor wrinkles in all trunk zones and of major wrinkles in the base and tip of the trunk. Adult Asian elephants have on average more major wrinkles in the shaft of the trunk (x_J= 115, SD = 26) than adult African elephants (x_J= 70, SD = 13; two-sample t-test (12) = 4, p = 0.001, d = 2.18).

**Figure S2:**
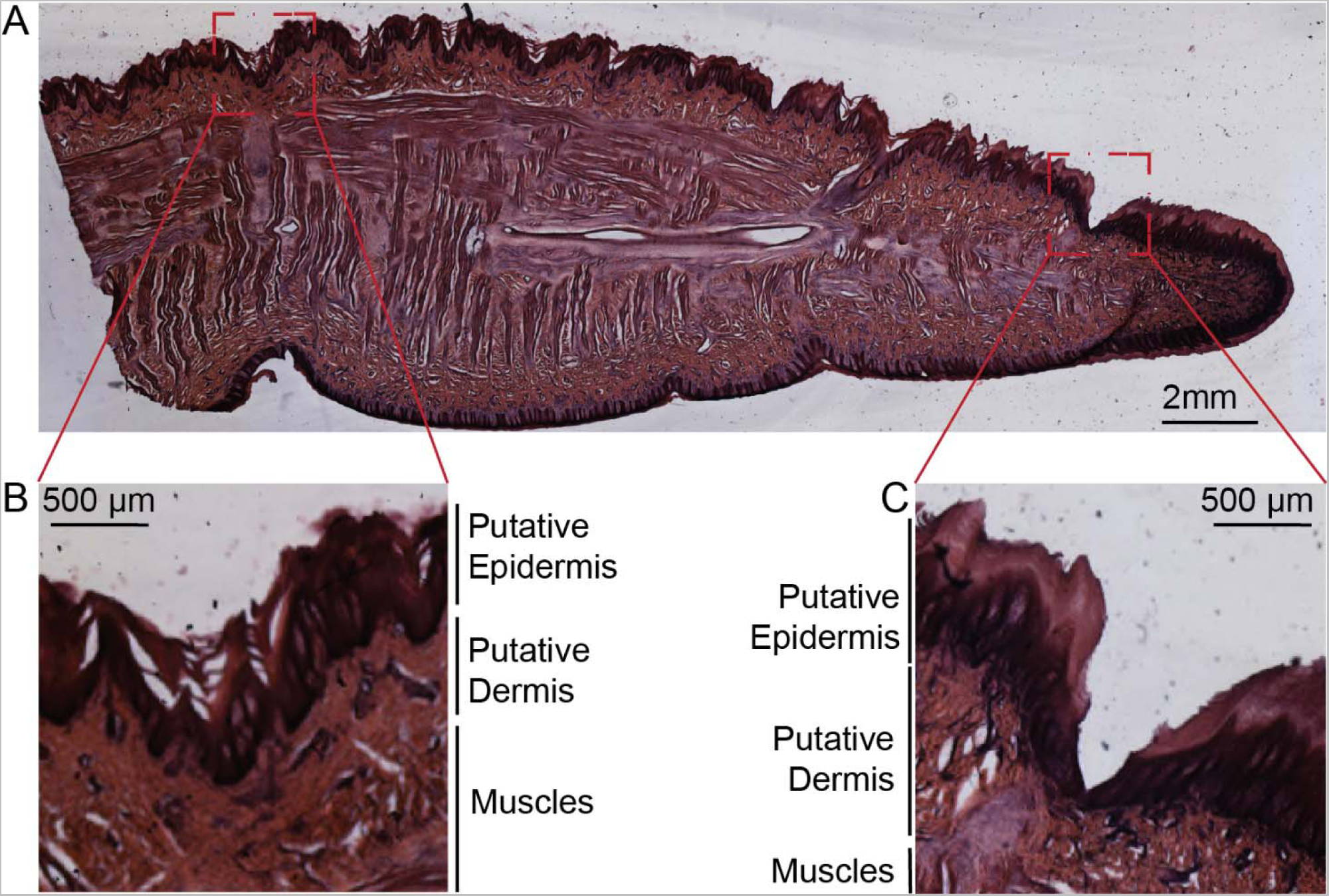
Hematoxylin-eosin staining of an Asian calf’s trunk tip. **A**, Parasagittal slice of the trunk tip of an Asian baby elephant, hematoxylin-eosin stained. Staining originally done for Deiringer et al. but not published **B**, High magnification view of proximal dorsal tip wrinkles in the Asian baby elephant trunk. The different skin layers visible with the staining protocol are labeled, including the putative epidermis, putative dermis, and transition to the muscles. **C**, High magnification view of distal dorsal tip wrinkles in the Asian baby elephant trunk. The different skin layers are visible with the staining protocol are labeled similarly to (B).

**Table S1:** Overview of elephant lab samples, treatment of specimen, and derived data.

| **Name**<br>**(species)** | **Sex** | **Location at**<br>**death** | **Age (y), died**<br>**on**<br>**(dd/mm/yy)** | **Specimen**<br>**treatment** | **Data derived** |
| --- | --- | --- | --- | --- | --- |
| Ilona (*E. maximus*) | F | Zoo Karlsruhe, Germany | ~45, 31/03/14 | Frozen for several years, in fixative (4% PFA) | Wrinkle numbers trunk tip |
| Vilja (*E. maximus*) | F | Wilhelma Stuttgart, Germany | ~61, 10/07/10 | Frozen for several years | Wrinkle numbers trunk tip |
| Naing Thein (*E. maximus*) | M | Zoo Copenhagen, Denmark | 40, 10/02/21 | In fixative (4% PFA) | Wrinkle numbers trunk tip |
| Asian Baby (*E. maximus*) | Unknown | Tierpark Hagenbeck, Germany | 0, 2012 | Frozen for several years, in fixative (4% PFA) | Wrinkle numbers trunk tip, shaft and base |
| Hoa's Baby (*E. maximus*) | F | Zoo Leipzig, Germany | 0, 2015 | Frozen for several years, in fixative (4% PFA), iodine stained for CT scanning | Wrinkle numbers trunk tip, shaft and most of base, CT scanned |
| Indra (*L. africana*) | F | Elefantenhof Platschow, Germany | ~35, 06/09/22 | In fixative (4% PFA) | Wrinkles numbers trunk tip, shaft, and base |
| Linda (*L. africana*) | F | Zoo Poznan, Poland | ~35, 02/02/21 | Frozen for several years | Wrinkle numbers trunk tip |
| Zimba (*L. africana*) | F | Opel-Zoo Kronberg, Germany | ~39, 10/04/21 | In fixative (4% PFA) | Wrinkle numbers trunk tip |
| Ali (*L. africana*) | M | Opel-Zoo Kronberg, Germany | ~24, 19/01/04 | Frozen for several years | Wrinkle numbers trunk tip |
| AM 1 (*L. africana*) | M | Colchester Zoo, Great Britain | 0, 2011 | Frozen for several years, in fixative (4% PFA), iodine stained for CT scanning | Wrinkle numbers trunk tip, shaft, and base, CT scanned |

**Table S2:** Overview of zoo elephants, photographs used, and derived data.

| **Name (*species*)** | **Sex** | **Location at time of photographs** | **Age at the time of photographs** | **Data derived** |
| --- | --- | --- | --- | --- |
| Anchali (*E. maximus*) | F | Zoo Berlin, Germany | 3 d; 10 y | Wrinkle numbers trunk tip, shaft, and base for baby and adult |
| Carla (*E. maximus*) | F | Zoo Berlin, Germany | ~49 y | Wrinkle numbers trunk tip, shaft, and base |
| Drumbo / Mary (*E. maximus*) | F | Zoo Berlin, Germany | ~53 y | Wrinkle numbers trunk tip, shaft, and base |
| Malka (*E. maximus*) | F | Zoo Tbilisi, Georgia | ~26 y | Wrinkle numbers trunk tip, shaft, and base |
| Pang Pha (*E. maximus*) | F | Zoo Berlin, Germany | 36 y | Wrinkle numbers trunk tip, shaft, and base |
| Elbrus / Grand (*E. maximus*) | M | Zoo Tbilisi, Georgia | 27 y | Wrinkle numbers trunk tip, shaft, and base |
| Victor (*E. maximus*) | M | Zoo Berlin, Germany | 29 y | Wrinkle numbers trunk tip, shaft, and base |
| Hoa's Baby (*E. maximus*) | F | Zoo Leipzig, Germany | 6 days old | Wrinkle number trunk base |
| Drumbo (*L. africana*) | F | Zoo Schönbrunn, Austria | ~46 y | Wrinkle numbers trunk tip, shaft, and base |
| Iqhwa (*L. africana*) | F | Zoo Schönbrunn, Austria | 9 y | Wrinkle numbers trunk tip, shaft, and base |
| Linda (*L. africana*) | F | Zoo Poznan, Poland | ~26 y | Wrinkle numbers trunk tip, shaft, and base |
| Mongu (*L. africana*) | F | Zoo Schönbrunn, Austria | 19 y | Wrinkle numbers trunk tip, shaft, and base |
| Numbi (*L. africana*) | F | Zoo Schönbrunn, Austria | ~28 y | Wrinkle numbers trunk tip, shaft, and base |
| Tonga (*L. africana*) | F | Zoo Schönbrunn, Austria | ~38 y | Wrinkle numbers trunk tip, shaft, and base |
| Kibali (*L. africana*) | F | Zoo Schönbrunn, Austria | 1 y | Wrinkle numbers trunk tip, shaft, and base |

